# Hingepoints and neural folds reveal conserved features of primary neurulation in the zebrafish forebrain

**DOI:** 10.1101/738161

**Authors:** Jonathan M. Werner, Maraki Y. Negesse, Dominique L. Brooks, Allyson R. Caldwell, Jafira M. Johnson, Rachel M. Brewster

**Affiliations:** Department of Biological Sciences, University of Maryland Baltimore County, Baltimore, MD 21250

## Abstract

Primary neurulation is the process by which the neural tube, the central nervous system precursor, is formed from the neural plate. Incomplete neural tube closure occurs frequently, yet underlying causes remain poorly understood. Developmental studies in amniotes and amphibians have identified hingepoint and neural fold formation as key morphogenetic events and hallmarks of primary neurulation, the disruption of which causes neural tube defects. In contrast, the mode of neurulation in teleosts such as zebrafish has remained highly debated. Teleosts are thought to have evolved a unique pattern of neurulation, whereby the neural plate infolds in absence of hingepoints and neural folds (NFs), at least in the hindbrain/trunk where it has been studied. We report here on zebrafish forebrain morphogenesis where we identify these morphological landmarks. Our findings reveal a deeper level of conservation of neurulation than previously recognized and establish the zebrafish as a model to understand human neural tube development.

## INTRODUCTION

Primary neurulation is the process by which the neural tube, the precursor of the brain and spinal cord, is shaped from the neural plate. These morphogenetic events have mostly been studied in amniotes (mouse and chick) and amphibians (frogs), where conserved mechanisms were identified. Following neural induction, the neural plate (Supplemental Figure S1A) narrows and elongates by convergent extension movements^1,2^. The morphology of the neural plate is further changed in a biphasic manner^3^. The first morphological event is the formation of the medial hingepoint (MHP), which bends the flat neural plate into a V shape, forming the neural groove (Supplemental Figure S1B). Neural Folds (NFs) at the edges of the neural plate subsequently elevate, a process that is further driven by head mesenchyme expansion^4^. In the second phase, the neural plate folds at paired dorso-lateral hingepoints (DLHPs), bringing the NFs closer together (Supplemental Figure S1C). The NFs eventually meet and fuse at the midline, completing neural tube formation. The dorsal midline is subsequently remodeled to separate the inner neuroectoderm from the outer non-neural ectoderm fated to become the epidermis (Supplemental Figure S1D). In the cranial region of amniotes, neural tube closure is initiated at several sites and extends via a zippering process from these closure points to seal the neural tube. Incomplete cranial neural tube closure occurs frequently, resulting in anencephaly and exencephaly^3^.

Several cellular mechanisms have been identified that contribute to the bending of the neural plate, among which apical constriction is the most studied^5–8^. A meshwork of F-actin accumulates at the apical cortex of neuroectodermal cells and contracts, thereby reducing the cell apex (Supplemental Figure S1b1). The contraction is driven by myosin that co-localizes with F-actin at the apical pole. Disruption of F-actin using drug inhibitors or genetic tools causes severe cranial neural tube defects^5,9–14^. Similarly, treatment with blebbistatin, an inhibitor of non-muscle myosin II activity, impairs apical constriction in the superficial layer of the *Xenopus* neural plate^15^.

In contrast to hingepoint formation, the cell intrinsic mechanisms that shape NF cells are less understood. The NFs are bilaminar, consisting of a layer of neuroectoderm capped by a layer of non-neural ectoderm^16^, a topology that is acquired via epithelial ridging, kinking, delamination and apposition^16,17^. NF fusion involves the formation of cellular protrusions that span the midline gap and establish the first contact and attachment points with NF cells from the contralateral side^18–20^ (Supplemental Figure S1c1). There is little consensus on the cell type (neuroectoderm or non-neural ectoderm) that generates the cellular protrusions as it varies depending on the species and the axial level. In the mouse forebrain, the initial contact is made by neuroectodermal cells^18,21–24^. Disruption of F-actin blocks the formation of protrusions and prevents NF fusion^20^, revealing the central role of these cellular processes.

While more anterior regions of the neural tube undergo primary neurulation as described above, most vertebrates also exhibit a distinct type of neural tube formation in more posterior regions, termed secondary neurulation. During this process, a mesenchymal cell population condenses into a solid neural rod that subsequently epithelializes and forms a central lumen^25,26^. In zebrafish, the neural tube in the hindbrain and trunk region initially forms a solid rod that later develops a lumen, a process seemingly analogous to secondary neurulation^27^. However, examination of the tissue architecture in zebrafish^28–30^ and other teleosts^31,32^ revealed that the neural rod is shaped by infolding of a neural plate (albeit incompletely epithelialized), which best fits the description of primary neurulation^33^. Despite this evidence, differences in tissue architecture, the multi-layered architecture of the neural plate and the apparent lack of hingepoints, neural groove and NFs are difficult to reconcile with a mode of primary neurulation and have contributed to the persistent view that neural tube formation in teleosts is different than in other vertebrates^34–36^.

We show here that, in contrast to the zebrafish hindbrain and trunk region, the process of neural tube formation in the forebrain exhibits many of the hallmarks of primary neurulation. We observe the presence of hingepoints and NFs in the epithelialized anterior neural plate (ANP), demonstrate that formation of the MHP involves oscillatory apical contractions that progressively reduce the apical surface, and show that disruption of myosin function impairs apical constriction of these cells and NF convergence. We further show that neural tube closure is initiated at two separate sites in the forebrain and that fusion of the NFs is mediated by filopodial-like extensions of neuroectodermal cells that bridge the midline. These findings identify conserved mechanisms of primary neurulation that were previously overlooked in teleosts and support the suitability of zebrafish for understanding the etiology of human neural tube defects.

## RESULTS

### Precocious epithelialization of the anterior neural plate is associated with bending

The zebrafish ANP is quite distinct from the neural plate in more posterior regions, as it undergoes precocious epithelialization^37^. To assess whether epithelialization correlates with a change in the mode of neurulation, we examined the morphology of the neuroectoderm in optical cross sections at developmental stages ranging from 2 to 10 somites (som). We observed that at 2-5 som the ANP has a V shape marked by a medial neural groove (black arrowhead) flanked by the elevated lateral edges of the ANP (white arrowhead), which are reminiscent of NFs (Figure 1A-B). By 7 som the groove is no longer visible and the elevated edges of the ANP have fused medially (Figure 1C, white arrowhead) and form a dorsal bulge (Figure 1C-D, white arrowhead). These observations suggest that the ANP bends at the midline, which correlates with elevation of the lateral edges of the ANP.

**Figure 1.**
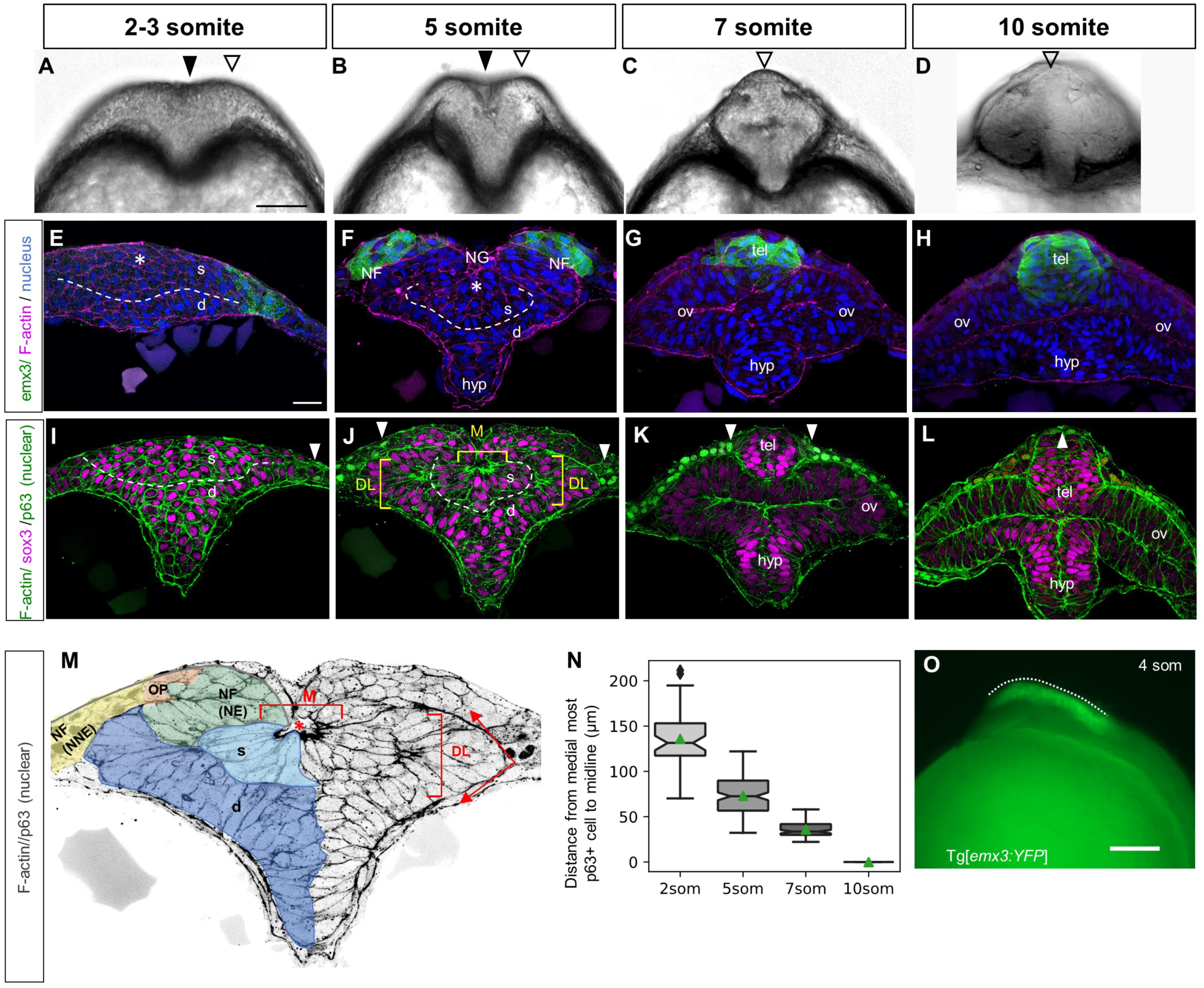
Hingepoints and neural folds contribute to forebrain morphogenesis. (A-D) Optical sections at the level of the forebrain of WT embryos at the 2-3 som (A), 5 som (B), 7 som (C) and 10 som (D) stages. (E-L) Transverse sections through the ANP of 2-3 som(E,I), 5 som (F,J), 7 som (G, K) and 10 som (H, L) embryos. (E-H) Tg[*emx3:YFP*] embryos labeled with anti-GFP (green), phalloidin (F-actin, magenta) and DAPI (nuclei, blue). (I-L) WT embryos labeled with phalloidin (F-actin, green), anti-Sox3 (magenta) and anti-p63 (nuclear label, green). (M) Higher magnification image of panel J, grey scaled to reveal F-actin and p63 and pseudo-colored - color code: light blue: superficial eye field cells, dark blue: deep layer cells that apically constrict to form the optic vesicles, green: neural component of the neural fold, orange: olfactory placode (Sox3/p63-negative cells), yellow: non-neural component of the neural fold. (N) Measurements of neural fold convergence, scored as the distance between the medial-most p63-positive cells on either side of the midline at different developmental stages. Notches depict the 95% confidence interval around the median and the green triangle depicts the distribution mean. 2 som: 48 measurements from 14 embryos, mean= 136; 5 som: 144 measurements from 16 embryos, mean=73.0; 7 som: 87 measurements from 10 embryos, mean=36.0; 10 som: p63 domain is fused, no measurements. Statistical analysis: Mann-Whitney U tests, two-sided; 2 som vs 5 som : P = 1.18e^-21; 2 som vs 7 som : P = 8.35e^-22; 5 som vs 7 som : P = 2.14e^-31. (O) Side view of a 4 som Tg[*emx3:YFP)* embryo. Abbreviations: s = superficial layer; d = deep layer; NF = neural fold; NG = neural groove; hyp = hypothalamus; tel = telencephalon; ov = optic vesicle; DL = Dorso-lateral hingepoints; M = Medial hingepoint; OP = olfactory placode; NE = neural ectoderm; NNE = non-neural ectoderm. Annotations: black arrowhead = median groove, white open arrowhead = elevated neural fold-like structure, dashed line = separation of the deep and superficial layers; brackets = hingepoints; dotted line = A-P range of the neural folds; white arrowhead = medial-most epidermis; red asterisk = neuropore. Scale bars: A and O= 100 μm, E= 25 μm.

### The anterior neural plate is multi-layered and gives rise to the eyes and forebrain

The central part of the ANP is fated to become the eyes, while its lateral edges produce the dorsally-located telencephalon and the ventral-most ANP region gives rise to the hypothalamus/ventral diencephalon^37,38^. To gain a better understanding of the morphogenetic events that shape the ANP, we examined transverse views of transgenic Tg[*emx3:YFP*] embryos at developmental stages 2, 5, 7 and 10 som, in which telencephalon precursors are labeled^39^. We observed, as previously described^37^, that the ANP is a multi-layered tissue, with a mesenchymal superficial core and a marginal or deep layer. These layers will henceforth be referred to as the superficial and deep layers, respectively (s and d in Figure 1E-F). The YFP-positive lateral edges of the ANP elevate, migrate over the eye field and fuse medially at 7 som to form the telencephalon (Figure 1G-H). By 7 som the multilayered ANP resolves into a single-cell layered neuroectoderm (Figure 1G). While the morphological changes and cellular dynamics that form the eyes are quite well understood^37,38^, the accompanying events that shape the forebrain are for the most part unknown and the focus of the current study.

### Presence of hingepoints and neural folds in the anterior neural plate

To investigate whether the ANP bends and folds around hingepoints to bring NFs in close apposition, we examined the cytoarchitecture of this tissue in embryos at stages 2-10 som, labeled with phalloidin (filamentous (F) actin), anti-Sox3 (neural cells) and anti-p63 (epidermal cells, nuclear label) (Figure 1I-L).

We found that at 2 som, neural cells appear mesenchymal with no visible polarized enrichment of F-actin (Figure 1I), consistent with previous observations^37^. The epidermis at this stage is in a far lateral position (white arrowhead in Figure 1I).

Between 2 and 5 som, clusters of cells in the superficial and deep layers of the eye field have been reported to undergo early epithelialization^37^, which we confirm here with foci of F-actin enrichment in the medial/superficial region (M in Figure 1J, M) and in two dorso-lateral clusters in the deep marginal layer (DL in Figure 1J, M). The apical surfaces of the superficial cells appear to constrict and orient towards the midline, resulting in the formation of a medial neural groove (asterisk in Figure 1M). Similarly, the paired dorso-lateral clusters of cells in the deep layer are also apically constricted and the neuroectoderm folds sharply at this level (red arrows in Figure 1M). These data suggest that the medial and dorso-lateral cells enriched for apical F-actin may function as hingepoints. Similarly to amphibians whose neural plate is bilayered^40^, the putative medial hingepoint in the zebrafish ANP forms in the superficial layer and is therefore more dorsally positioned than its chick and mouse counterpart. However, these cells eventually sink inwards, as revealed by fate mapping using photoconvertible membrane-targeted Kaede (mKaede) to label this cell population (Supplemental Movie 1). These findings are consistent with a previous study showing that superficial cells radially intercalate between deep marginal cells^37^, which contributes to the expansion of the optic vesicles (Figure 1G-H). Zebrafish dorso-lateral hingepoints do not undergo these cellular rearrangements as they form in the deep layer.

The lateral edges of the ANP are bilaminar at 5 som, consisting of a layer of neuroectoderm cells capped by a layer of p63-positive non-neural ectoderm (and Sox3/p63-negative olfactory placodal cells bridging the two layers), indicative of a NF structure (Figure 1J, M). The YFP-positive ANP cells in Tg[*emx3:YFP]* embryos correspond to the neuroectoderm component of the NF (Figure 1F), revealing that the tip of the NF gives rise to the telencephalon. At 4 som, the YFP-positive region of Tg[*emx3:YFP*] embryos extends the length of the forebrain (Figure 1O), delineating the anterior-posterior range of the NFs.

The NF and putative hingepoints are transient as they are no longer observed in 7 som embryos. By this stage, the tips of the NFs have converged medially and fused, forming the telencephalon (Figure 1K). These cells are enriched in apical F-actin at 10 som, suggesting that they epithelialize (Figure 1L). The non-neural ectoderm still occupies a lateral position at 7 som (Figure 1K, arrowheads), however by 10 som these cells migrate and fuse dorsally (single arrowhead in Figure 1L), indicating that, as observed in mice, the neuroectodermal component of the NFs meet first (Supplemental Figure S1)^21,22^. Measurements of the distance between the medial-most p63-positive domain and the dorsal midline at stages 2-7 som indicate that the non-neural ectoderm portion of the NFs converges steadily towards the midline and may provide a lateral force that contributes to the displacement of NFs (Figure 1N).

These observations reveal that transient medial and dorso-lateral epithelialized cell clusters may be the functional equivalents of the MHP and paired DLHPs of amniotes (and will be referred to henceforth as such), as they form at the right time and place to contribute to the formation of the neural groove (MHP), bending of the neuroectoderm and medial convergence of the NFs.

### Cell shape changes underlying dorso-lateral hingepoint and neural fold formation

To capture the dynamics of NF formation and image cells at higher resolution, we mosaically expressed membrane-targeted GFP (mGFP) and imaged embryos at the 2-10 som stages in transverse sections (Figure 2). 3D reconstructions of some of these images were generated to visualize the spatial relation between labeled cells (Supplemental Movie 2, 3).

**Figure 2.**
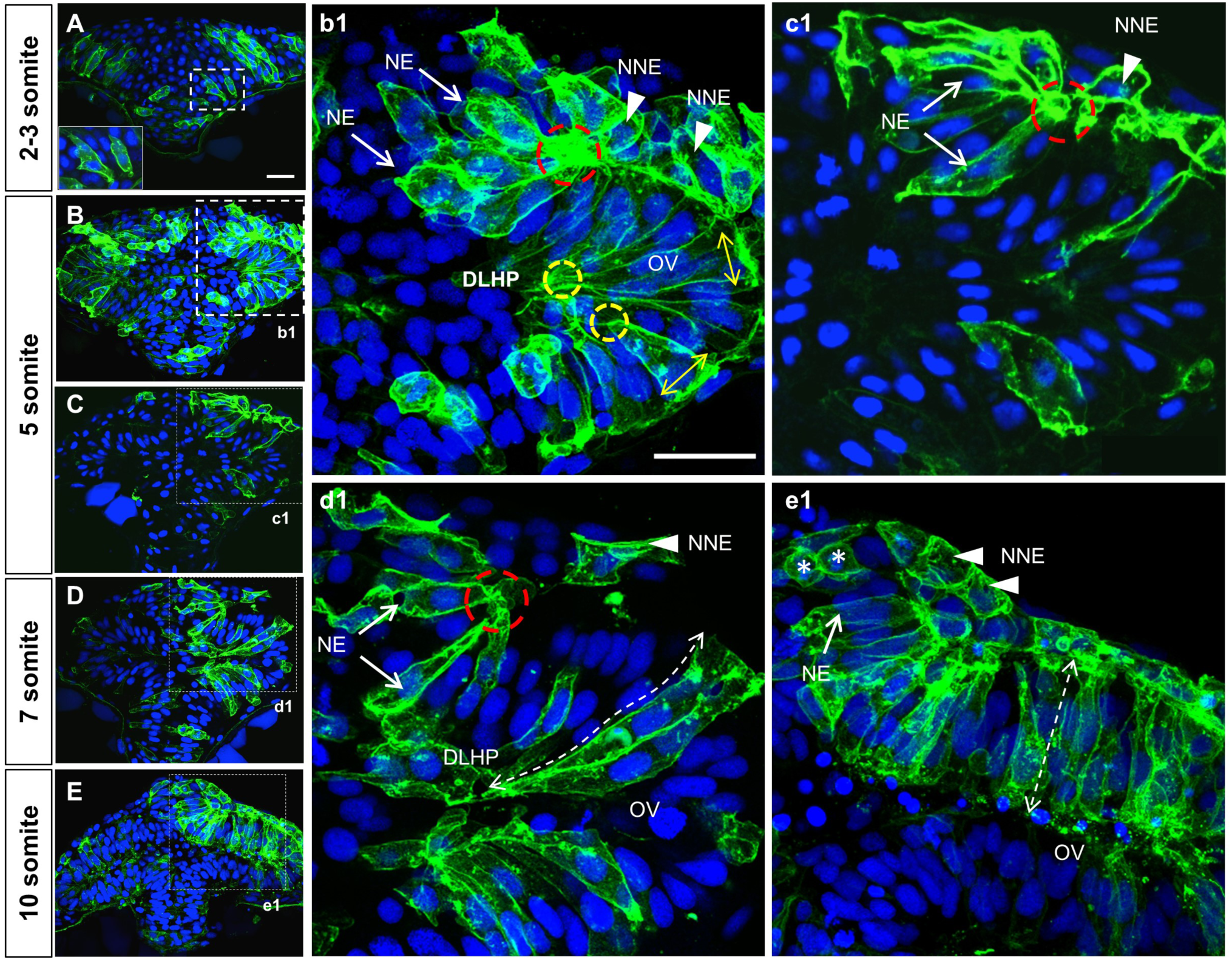
Cell shape changes in the deep layer of the ANP that contribute to DLHP and neural fold formation. (A-E) Transverse section, at the level of the forebrain, of embryos at the 2-3 (A), 5 (B, b1, C, c1), 7 (D, d1) and 10 (E, e1) somite stages mosaically-expressing mGFP (green) and labeled with the nuclear marker DAPI (blue). The inset in A is a higher magnification of dashed area in A. (b1-e1) Higher magnifications of regions delineated by dotted lines in (A-E). Annotations: red dashed circle = basal constriction of NE component of neural fold; yellow circle = apical constriction of DLHP cells; arrows = NE component of neural fold; arrowheads = NNE component of neural folds; double dashed arrow = elongated deep cells of the optic vesicle, asterisks = dividing cells in the prospective telencephalon. Scale bars: 25 μm in A and b1.

At 2 som, cells in the deep layer have a columnar shape with one end in contact with the basal lamina and a future apical surface oriented towards the midline (Figure 2A). These cells extend membrane protrusions into the superficial layer, which may promote radial intercalation (inset in Figure 2A).

At 5 som, cells in the dorso-lateral deep layer have undergone apical constriction and basal expansion, forming DLHPs (yellow dotted circles and double arrowheads in Figure 2B,b1; Supplemental Movie 2). These cell shape changes may initiate the outpocketing of the optic vesicles (ov in Figure 2B,b1). The bilaminar organization of the NFs is clearly visible at this stage. Neuroectodermal cells within the NFs (arrows in Figure 2b1 and c1) are elongated and their basal poles are constricted, giving them the appearance of fanning out from a focal point (red circles in Figure 2b1 and c1). Furthermore, their plasma membrane is ruffled, indicative of protrusive activity (Supplemental Movie 2).

By 7 som (Figure 2D, Supplemental Movie 3), neuroectodermal cells of the NFs have reached the dorsal midline. They maintain their organization with basal poles constricted and clustered at a focal point on the basement membrane (red circle in Figure 2d1). 3D reconstructions reveal that these cells are finger-shaped as they extend across the dorsal midline. DLHP cells maintain their apical constriction/basal expansion and further elongate, contributing to the expansion of the optic vesicles (dotted double arrowhead in Figure 2d1).

At 10 som (Figure 2E, e1) deep cells in the eye field and NFs have a columnar, epithelial organization. Consistent with previous findings, eye field cells shorten along their apico-basal axis by contracting their apical processes and coincidently transition to a more dorso-ventral orientation^37^ (dotted double arrow in Figure 2e1). Cuboidal non-neural ectoderm cells cover the dorsal surface of the newly formed telencephalon (arrowheads in Figure 2e1.

The dynamic cell shape changes in the deep layer of the neuroectoderm were quantified by measuring cell length and the apico:basal ratio of mGFP-labeled cells located at different positions along the medio-lateral axis of the ANP at 2, 5 and 7 som (Figure 3). These data reveal that the average length of DLHP cells increases while their apical:basal ratio decreases between 2 and 7 som, coincident with optic vesicle evagination. They also confirm that neuroectodermal cells of the NFs adopt the opposite configuration (reverse hingepoints), with basally constricted poles.

**Figure 3.**
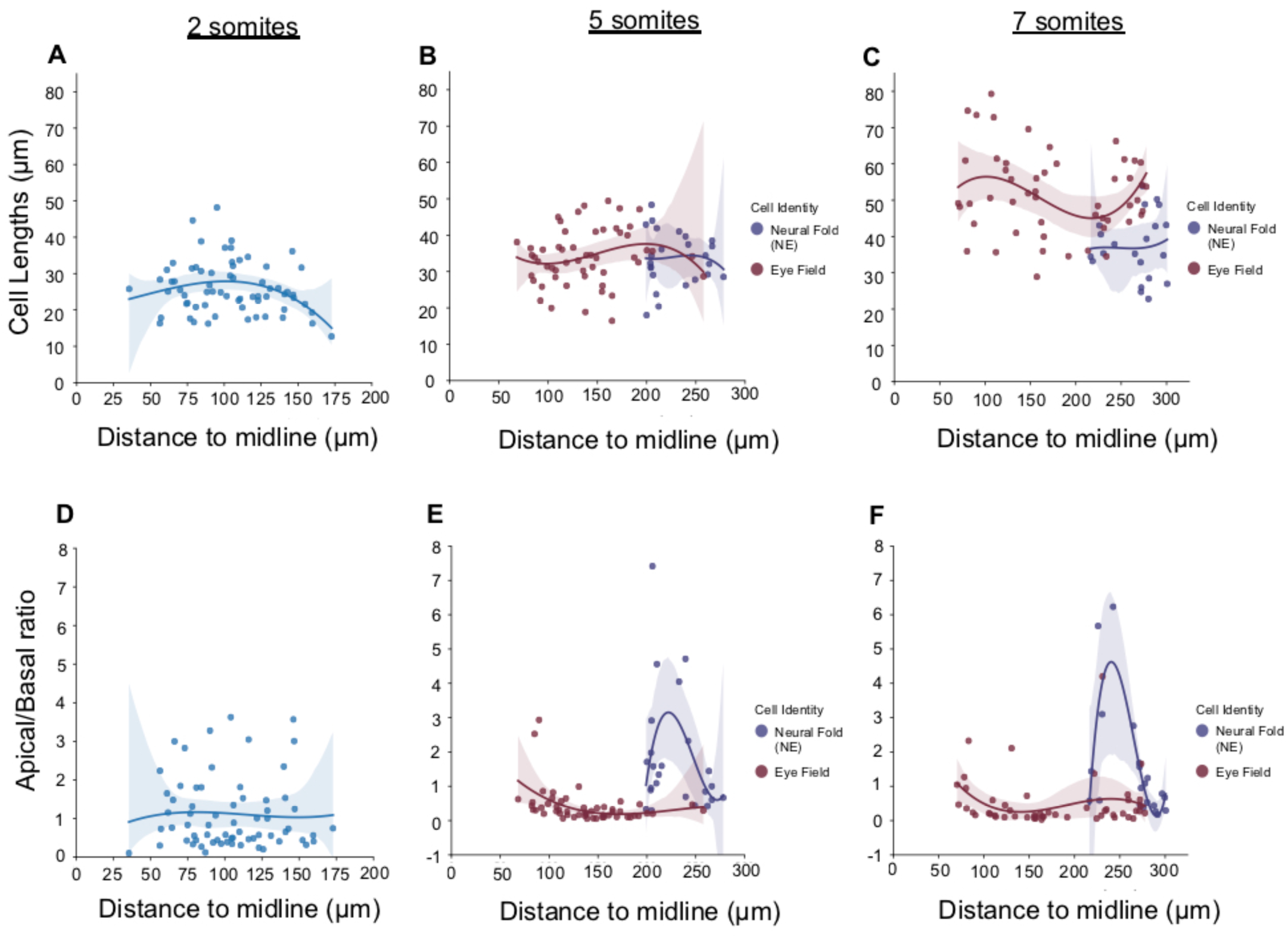
Measurements of cell shape changes in the deep neuroectodermal layer and neural folds. (A-C) Measurements of cell lengths (μM, Y axis) of m-GFP-labeled cells at different positions relative to the ANP midline (μM, X axis). Measurements begin in the region fated to become the optic vesicles. (D-F) Measurements of the apico:basal surface ratio of m-GFP-labeled cells at different positions relative to the ANP midline (μM, X axis). Measurements begin in the region fated to form the optic vesicles. The original scatter plot was fitted to a 3rd degree polynomial. Confidence intervals for the polynomial are 95% and were calculated with bootstrap sampling (n=1000). Color code: blue = neuroectodermal cells of the deep layer of 2 som stage embryos that are not yet identifiable based on cellular morphology; red = cells that form the optic vesicles; blue = cells that form the neuroectodermal component of the neural folds.

To capture the dynamics of NF formation, we performed transverse view time-lapse imaging of WT embryos expressing mKaede or Tg[*emx3:YFP*] transgenic embryos expressing membrane-targeted RFP (mRFP) (n=2 embryos, Figure 4; Supplemental Movies 1 and 4. We observed that as the NFs elevate (blue dotted line in Figure 4A-C), NF cells basally constrict (red outline and arrow in Figure 4a1-c1). Interestingly, this cell behavior appears restricted to cells within the *emx3* expression domain (Figure 4D-f1).

**Figure 4.**
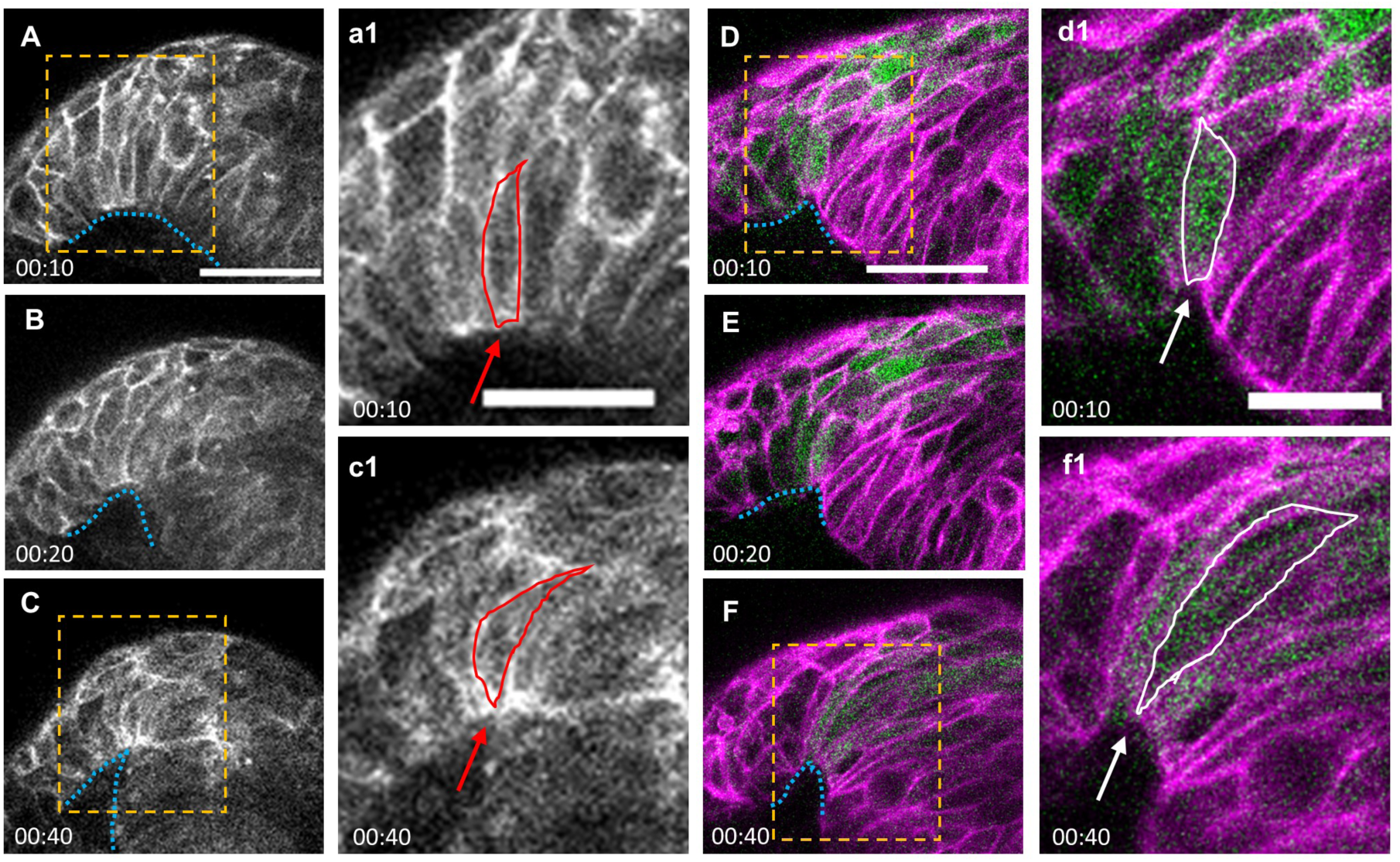
Dynamics of neural fold formation. A-F. Time lapse movie frames of the ANP, from a transverse view. A-C. Still frames of an embryo expressing membrane Kaede (mKaede). D-F. Still frames of a Tg[*emx3:YFP*] embryo expressing membrane RFP (mRFP, pseudo labeled magenta) and YFP (green). Yellow boxes in A, C, D and F identify magnified areas in a1, c1, d1 and f1. Annotations: blue dotted line: outlines the basal side of the neural folds; red and white lines identify individual neural fold cells; arrows: indicate narrowing surface in neural folds cells. Scale bars: 50 μm in A and D; 25 μm in a1 and d1.

These findings indicate that zebrafish DLHP cells adopt a wedge shape similar to their counterparts in amphibians and amniotes, which enables epithelial folding. They further identify basal constriction of NF cells as a cell shape change that contributes to NF formation.

### Medial hingepoint cells constrict apically and elongate

While mosaic expression of mGFP enabled high resolution imaging of DLHPs and NFs, it resulted in fewer cells labeled in the medial superficial layer, where the MHP forms. Phalloidin labeling was therefore used to image these cells in 2 and 5 som embryos.

At the 2 som stage, some medial/superficial cells immediately below the enveloping layer (EVL) are apically constricted, forming the MHP (Figure 5A-a2). By 5 som, MHP cells appear more densely packed, the majority of them are apically constricted and oriented towards the midline. Concomitant with these cell shape changes and the medial convergence of NFs, MHP cells shift to a more ventral position (Figure 5B-b2). A small opening, the neural groove, is observed immediately above the MHP at this stage (NG in Figure 5b1).

**Figure 5.**
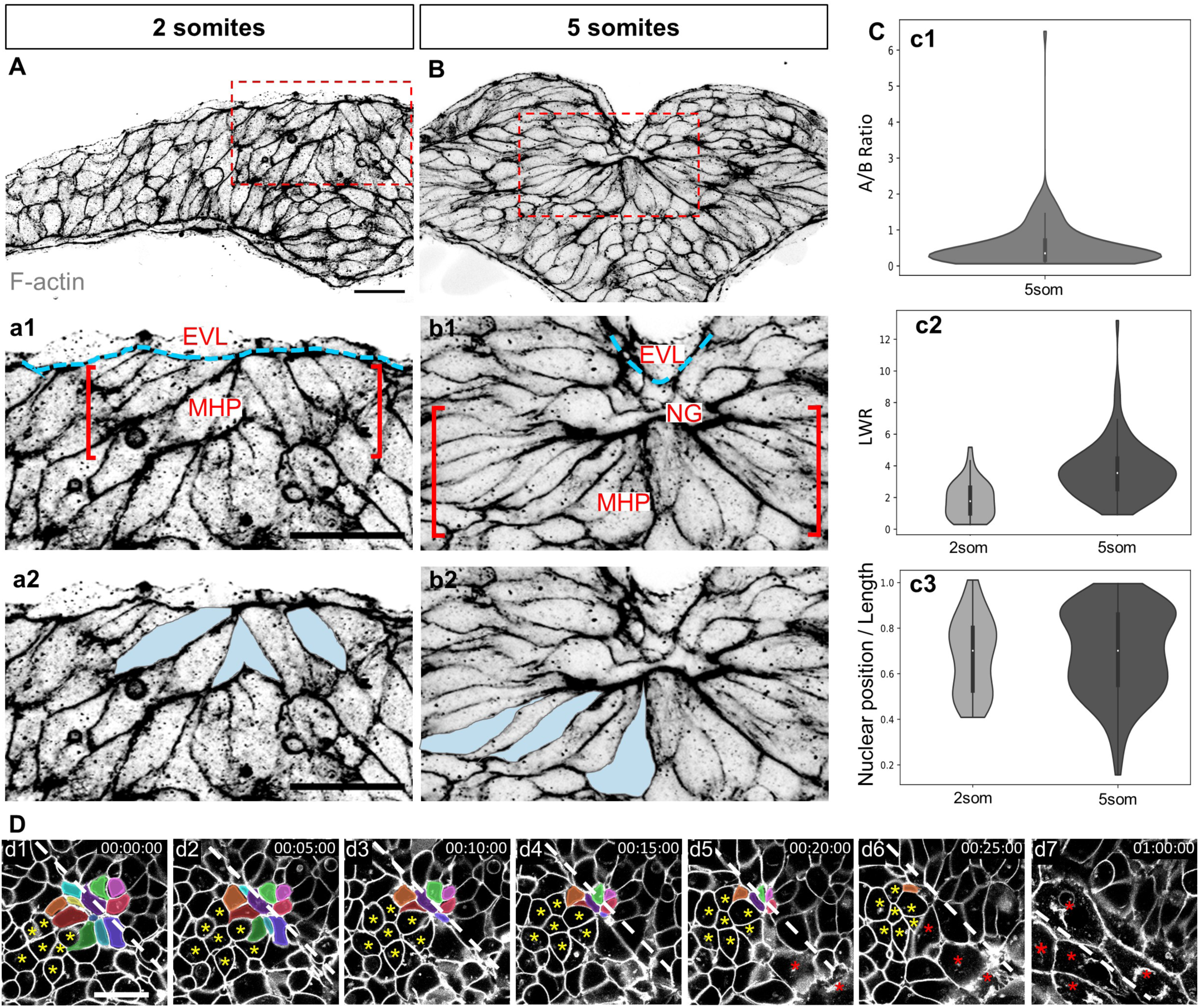
Apical constriction of MHP cells. (A-b2) Transverse sections through the ANP at the 2 (A, a1, a2) and 5 (B, b1, b2) som stages labeled with phalloidin (shown in greyscale). (a1-b2) are higher magnifications of the boxed areas in A and B, revealing the organization of the medial ANP (a1, b1) and the shape of individual MHP cells pseudo-colored in blue (a2, b2). (C) Quantitation of cell shape changes. Boxplot elements depict quartiles with the centerline depicting the median. (c1) Measurements of apical:basal surface ratio at 5 som (n= 115 cells from 4 embryos, mean=0.548). (c2) Measurement of length-to-width (LWR) ratio at 2 som (n= 47 cells from 5 embryos, mean=1.88) and 5 som (same cells as in c1, mean=3.70). A Mann-Whitney two-sided U Test revealed that the LWR increase between 2 som and 5 som is statistically significant (P = 7.80e^-11). (c3) Relative position of nucleus at 2 and 5 som measured in the same cell populations (c2). Mean nuclear position/cell length (0.682 at 2 som vs 0.696 at 5 som) is not statistically significant using a Mann Whitney U test (P=0.419). (D) Still frames of time lapse movie of m-GFP labeled embryo imaged from a dorsal view. Individual MHP cells are pseudo-colored, A cluster of cells adjacent to the MHP is indicated with yellow asterisks and EVL cells are labeled with red asterisks. Abbreviations: EVL = enveloping layer; MHP = medial hingepoint; NG = neural groove. Annotations: white dashed line= midline, red brackets = MHP region, blue dashed lines = outlines EVL, yellow asterisks: cells adjacent to MHP, red asterisks: EVL cells. Scale bars: 25 μm in A, a1 and a2; 10 μm in d1.

Measurements of the apico:basal surface ratio of MHP cells in 5 som embryos confirmed that they are wedge-shaped (Figure 5c1). Similar to amphibians^41^, apical constriction appears tightly coupled to cell elongation, as the length-to-width ratio (LWR) of MHP cells increases significantly between 2 and 5 som (Figure 5c2). Thus, both apical constriction and cell elongation appear to shape the MHP and contribute to tissue-level morphogenesis.

Another contributing factor to hingepoint formation in amniotes is the basal location of nuclei in both the medial and lateral hingepoints^42,43^. However, in zebrafish embryos, the relative position of the nucleus in medial superficial cells at 2 and 5 som does not change significantly and is therefore unlikely to contribute to wedging of these cells.

### Oscillatory constrictions reduce the apical surface of medial hingepoint cells

To gain a better understanding of the dynamics of apical constriction and cell internalization, the ANP of embryos ubiquitously expressing mGFP was imaged from a dorsal view using time-lapse microscopy between 2 som to 4 som (n = 2 embryos, Figure 5D and Supplemental Movie 5). The focal plane was set immediately below the EVL, at the level of the MHP. These movies revealed clusters of medially-located cells that undergo progressive cell surface reduction (color-coded in Figure 5d1-d7), while the surface area of adjacent, more lateral cells remained unchanged for the duration of imaging (yellow asterisks in Figure 5d1-d6). EVL cells came into focus immediately posterior to the cells with reducing apices. Since the EVL is drawn inward as a result of its close contact with apically constricting MHP cells (Figure 5B-b2), we surmise that apical constriction proceeds in a posterior-to-anterior direction. In later movie frames, EVL cells are no longer observed within the field of view, coinciding with the proximity of the NFs to the midline (Supplemental Movie 5). This indicates that medial EVL cells eventually lose contact with the MHP and return to their original position, allowing the NFs to fuse at the dorsal midline (Figure 1J-K).

To further examine the dynamics of apical constriction, the surface area of superficial ANP cells was measured at shorter intervals. Midline cells presumed to be part of the MHP (20 μm on either side of the midline, Figure 6A, E) or cells immediately adjacent to the MHP (greater than 20 μm on either side of the midline, Figure 6B) were scored. This analysis revealed that individual cells in both populations undergo oscillatory constrictions. Between constrictions, the surface areas of cells re-expand with gradually decreasing amplitude (Figure 6A, E), which is most pronounced in MHP cells. The time of oscillation between two expanded states revealed no significant difference between both groups (Figure 6C, median oscillation time of 45 seconds). Likewise, there was no significant difference in the timing of individual constrictions. However, it appears that MHP cells spent less time expanding than MHP-adjacent cells (15 seconds median time for MHP vs 30 second for MHP-adjacent, Figure 6D).

**Figure 6.**
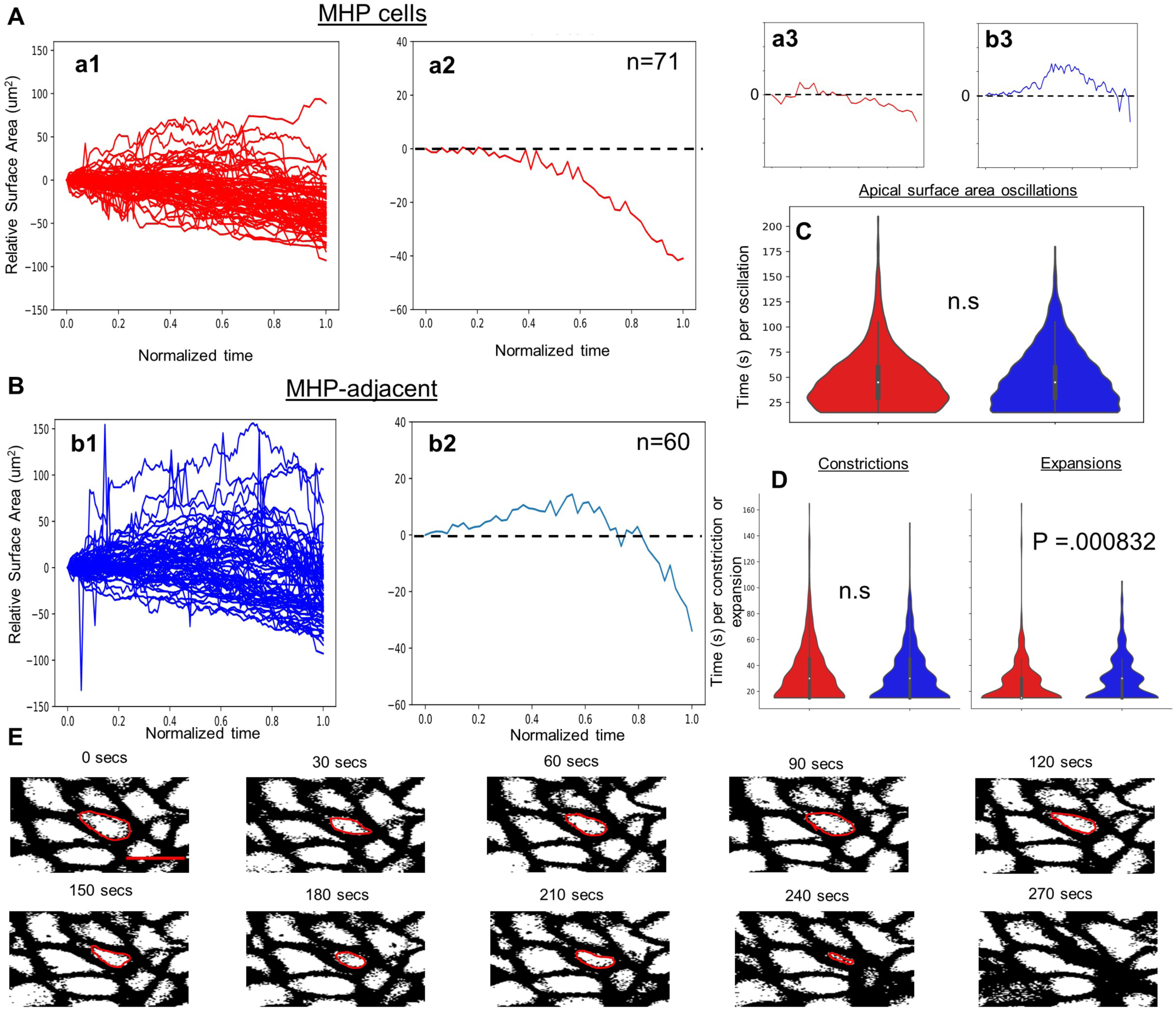
Oscillatory constriction with decreasing amplitude reduces the apical surface of MHP cells. (A) Measurements of medial, MHP cells. (a1) Relative apical surface areas over time for individual MHP cells. (a2) Median values of MHP relative apical surface areas over time, 95% confidence interval, n=71. (a3) Representative trace of relative apical surface area over time for an individual MHP cell. (B) Measurements of MHP-adjacent cells. (b1) Relative apical surface areas over time for individual MHP-adjacent cells. (b2) Median values of MHP-adjacent relative apical surface areas over time, 95% confidence interval. (b3) Representative trace of relative apical surface area over time for an individual MHP-adjacent cell. (C) Distributions of the duration of oscillation between two expanded states for MHP cells (red, median of 45 seconds per oscillation) and MHP-adjacent cells (blue, median of 45 seconds per oscillation). No significant difference (two-sided Mann Whitney U test, P=0.293, n=758 MHP cell oscillations, n=1,113 MHP-adjacent oscillations). (D) Distributions of the timing of apical constrictions or expansions for MHP cells (red) and MHP-adjacent cells (blue). There is no significant difference between the two groups for constriction time (two-sided Mann Whitney U test: constrictions, P=0.541, n= 458 MHP cell constrictions, n=640 MHP-adjacent constrictions) but there is a statistically significant difference for expansion time (P = 0.000832, n= 367 MHP-cell expansions, n=596 MHP-adjacent expansions). The median time for individual expansions of MHP and MHP-adjacent cells is 15 and 30 seconds respectively. All boxplot elements depict quartiles with the centerline depicting the median. (E) Still frames of time-lapse movie of m-GFP labeled cells shown in grey-scale. The oscillatory behavior of one cell, outlined in red, is shown over time. Scale bar: 10 μm in c1.

Together these data indicate that MHP cells and their neighbors undergo progressive narrowing of their apical pole via oscillatory constrictions and that these cellular dynamics proceed in a posterior-to-anterior direction.

### Neural fold fusion is initiated at closure points and mediated by protrusive activity

In mice, neurulation proceeds unevenly along the anterior-posterior axis with multiple closure initiation sites^3^, raising the question of whether NF fusion also occurs asynchronously in zebrafish. To address this, we performed time-lapse imaging of embryos mosaically expressing mGFP from a dorsal view around the time when opposing NFs approach the midline (n= 2 embryos, Figure 7A and Supplemental Movie 6). Neuroectodermal cells of the NF were identified based on their elongated shape and dorsal location.

**Figure 7.**
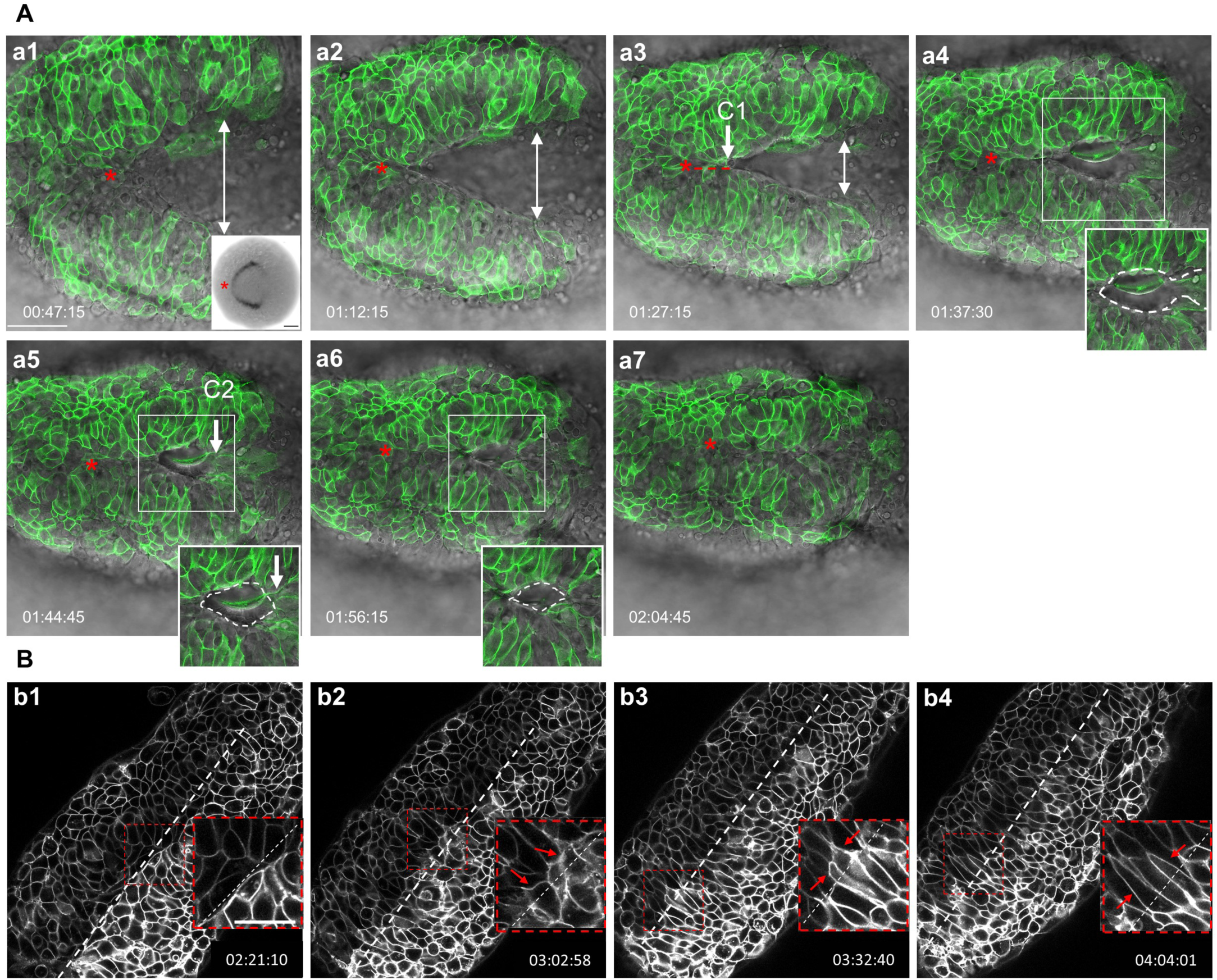
Dynamics of neural fold fusion. (A, a1-a7) Time lapse movie frames of an embryo expressing mosaic GFP, imaged from a dorsal view, showing the initiation of neural tube closure. Images are overlays of the green and brightfield channels. Inset in a1 shows a dorsal view of an *emx3*-labeled embryo. Insets in a4-a6 outline the eye-shaped opening that forms between closure sites one (CP1) and two (CP2) (B) Grey-scale time-lapse movie frames of an mGFP-labeled embryo imaged from a dorsal view, revealing the final stages of neural fold fusion. Insets in the lower right corner of panels b1-b4 are higher magnification views of boxed areas. Abbreviations: C1, C2: closure sites one and two. Annotations: red asterisk: apex of the neural fold arc; red dotted line: synchronous and posteriorly-directed neural fold fusion anterior to closure point one; white double arrows: distance between the neural folds; white dotted oval: eye shaped opening, the corners of which are defined by closure points one and two; white dotted line: embryonic midline; red arrows: filopodia extending across the midline; time-elapse is shown at the bottom of each panel. Scale bars: 50 μm in a1, 100 μm in a1 inset, and 25 μm in b1.

These movies revealed that the NFs have an arc shape with the apex positioned anteriorly (red asterisks in Figure 7A), similar to the expression domain of the telencephalon marker *emx3* at the onset of NF convergence (inset in Figure 7A). NF fusion is initiated near the apex of the arch and proceeds in an anterior-to-posterior direction for a distance of approximately 35 μm (red dotted line in Figure 7a3). At this level, defined as closure point one (C1 in Figure 7a3), NF fusion proceeds asynchronously as a second closure initiation site is formed more posteriorly (C2 in Figure 7A5). These closure points define an eye-shaped opening (white dotted oval in Figure 7a4, a5) and a zippering process begins at the anterior and posterior corners of this structure, progressing from both ends toward the middle (Figure 7a6,a7; Supplemental Movie 6).

The events that complete NF fusion in amniotes involve the extension of dynamic cellular projections towards the midline^18–20^. Likewise, we observed that zebrafish neuroectodermal cells extend filopodia to establish contact with cells from the contralateral side. Furthermore, the cell bodies appear to hyper-extend beyond the midline (Figure 7b3, b4 and Supplemental Movie 7), suggesting that they interdigitate between cells of the opposing NF. These cellular extensions are eventually retracted, as the midline becomes well defined after epithelialization (Figure 1L).

The presence of closure points and the usage of filopodia to establish contact with NF cells across the midline reveal additional aspects of forebrain neurulation that are conserved in zebrafish.

### Temporal overlap in the timing of key cell shape changes in the anterior neural plate

In amniotes, MHP formation precedes the shaping of paired DLHPs and initiates NF elevation. To reveal the relative timing of cell shape changes in zebrafish and their contributions to ANP morphogenesis, we plotted tissue dynamics using measurements from transverse time-lapse movies (n= 2 embryos, data shown for 1 embryo, Supplemental Movie 1). We observed that changes in the neural groove angle (indirect measurement of MHP apical constriction), the optic vesicle angle (which reflects DLHP apical constriction) and the basal NF angle (a proxy for basal constriction) occur concurrently and temporally correlate with NF elevation and convergence (Figure 8). This analysis revealed that, in contrast to amniotes, key cell shape changes in the ANP do not occur in a clear chronological manner and that one or more of these events are likely to contribute to ANP morphogenesis.

**Figure 8.**
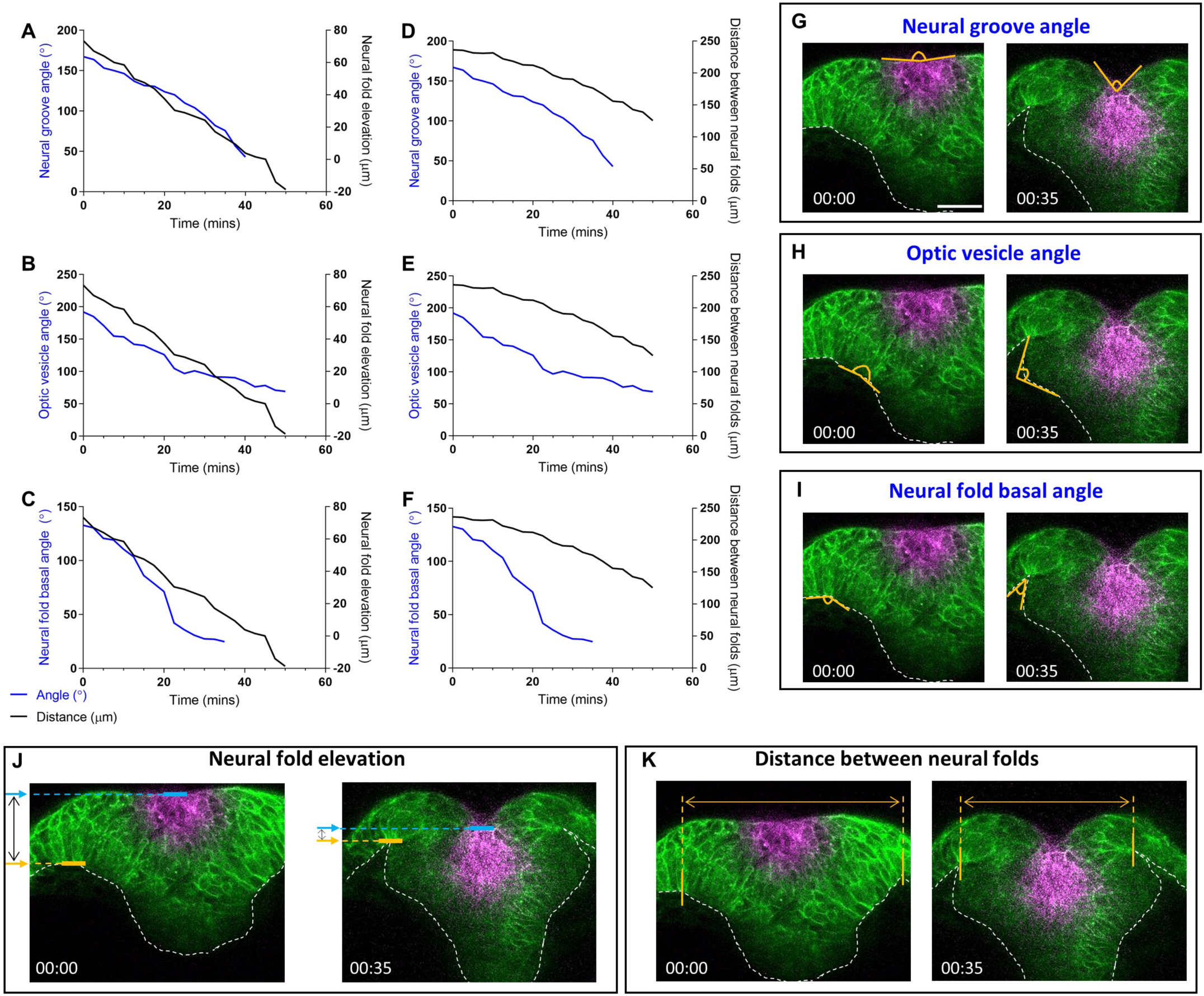
Dynamics of anterior neurulation. A-F. Graphs illustrating the dynamics of neural groove formation (left Y axis, blue line in A and D), optic vesicle angle (left Y axis, blue line in B and E) and neural fold basal angle (left Y axis, blue line in C and F) as compared to neural fold elevation (right Y axis, black line in A-C) and distance between the neural folds (right Y axis, black line in D-F) over time (X axis). G-K. Still frames of an embryo expressing mKaede (green), in which the MHP cells were photoconverted (magenta), showing how the measurements in graphs A-F were acquired, at two discrete time points. G. The neural groove was measured as the angle formed by the dorsal most tissue. H. The optic vesicle angle was measured as the angle formed by the outline of the optic vesicle as it forms. I. The neural fold basal angle was calculated as the angle formed by the basal side of neural fold cells. J. Neural fold elevation was measured as the distance between the basal side of neural fold cells and the apical side of MHP cells. As the neural folds elevate, that distance decreases and eventually becomes negative as the neural folds elevate above the MHP. K. Distance between the neural folds was measured as the distance between the basal side of neural fold cells. Scale bar: 50 μm.

### Molecular characteristics of medial and lateral hingepoints

Key features of cells that form hingepoints include apico-basal polarization as well as an accumulation of an apical contractile machinery composed of actin filaments (F-actin) and non-muscle myosin II^44^. To test whether the MHP and DLHPs in the zebrafish forebrain have some or all of these characteristics, the localization of apical markers Pard3-GFP (transiently expressed following mRNA injection), ZO-1 (anti-ZO-1) and N-cadherin (anti-N-cad) was examined along with F-actin (phalloidin) and phospho-Myosin Light Chain II (anti-P-MLC) in 5 som embryos (Figure 9).

**Figure 9.**
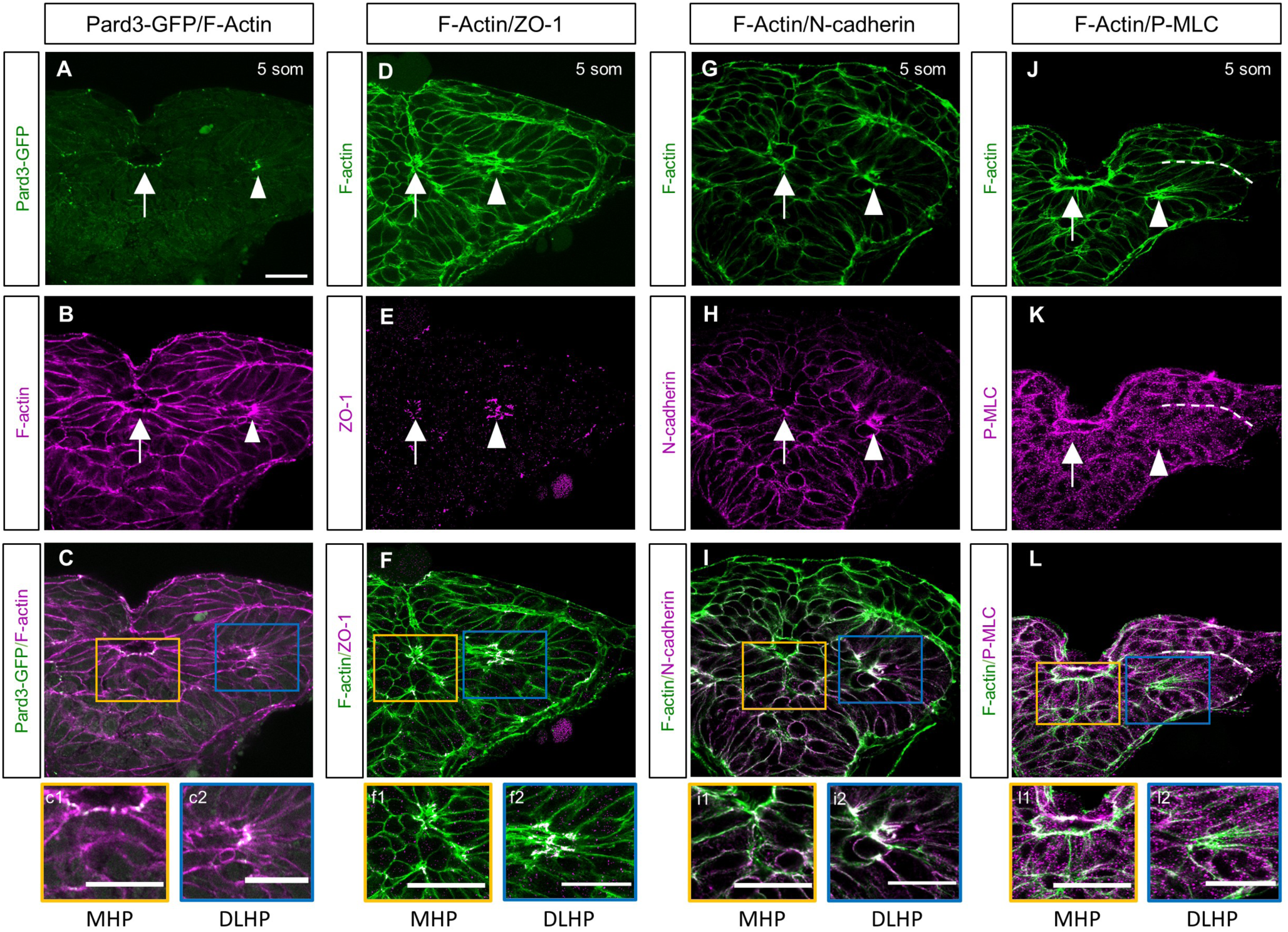
Molecular characterization of the MHP. (A-L). Transverse sections through the ANP at the 5som stage. Embryos double-labeled with Pard3-GFP (green) and phalloidin (F-actin, magenta) (A-c2); ZO-1 (magenta) and F-actin (green) (D-f2); N-cadherin (magenta) and F-actin (green) (G-i2); and with anti-P-MLC (magenta) and F-actin (green) (J-l2).(C, F, I, L) Magenta and green channel overlay. Insets show higher magnification images of the MHP (yellow box) and DLHP (blue box). Annotations: arrow = MHP; arrowhead = DLHP; dotted line (in J-L) = interface between the NE and NNE layers of the neural folds. Scale bars: 25 μm.

Apical co-localization of F-actin with Pard3-GFP (Figure 9A-c2), ZO-1 (Figure 9D-f2), and N-cad (Figure 9G-i2) was confirmed in both the MHP and DLHPs at 5 som. Thus, while the establishment of apico-basal polarity is generally delayed in the zebrafish neural plate relative to amniotes, the cell clusters that undergo apical constriction in the ANP epithelialize precociously. P-MLC accumulation at the apical pole is also apparent by the 5 som stage in MHP cells where it overlaps with F-actin (Figure 9J-l2). In contrast, P-MLC apical enrichment is not observed in the DLHPs (arrowhead in Figure 9K,l2). P-MLC does however accumulate in the cell cortex of all neuroectodermal cells where it overlaps with F-actin, in addition to the basal surface of EVL cells (arrow in Figure 9K, l1) and at the interface between the neuroectoderm and non-neural ectoderm (dotted line in Figure 9K-L).

These findings confirm that both the MHP and DLHP undergo early epithelialization and accumulate apical markers. However, the MHP and DLHPs are molecularly distinct structures given that the latter is not enriched for P-MLC.

### Myosin contractility is required for medial hingepoint formation and neural fold convergence

During neural tube closure, the actomyosin cytoskeleton is thought to be a driving force for apical constriction^41^. To address whether this molecular motor mediates apical constriction in zebrafish, non-muscle myosin II (NMII) was blocked using blebbistatin and a translation-blocking morpholino (MO) targeting NMIIb^45^ and the effect was analyzed in 5 som embryos that were labeled to reveal the localization of F-actin (phalloidin) and P-MLC (anti-P-MLC). Both treatments resulted in the absence of a clearly defined MHP (Figure 10a1,a1’ vs 10b1, b1’, c1 and c1’). LWR scores for these cells were close to 1 (Figure 10D), indicative of cell rounding, which prevented measurement of their apical:basal surface ratio. In contrast, while the length of DLHPs in treated embryos was reduced (Figure 10D), these cells retained their wedge shape (Figure 10a2,a2’ vs b2, b2’, c2, c2), with apical:basal surface ratios of less than 1 (Figure 10E). These observations are consistent with the lack of apical P-MLC enrichment in DLHP cells and support a conserved function for actomyosin in driving apical constriction of MHP cells.

**Figure 10.**
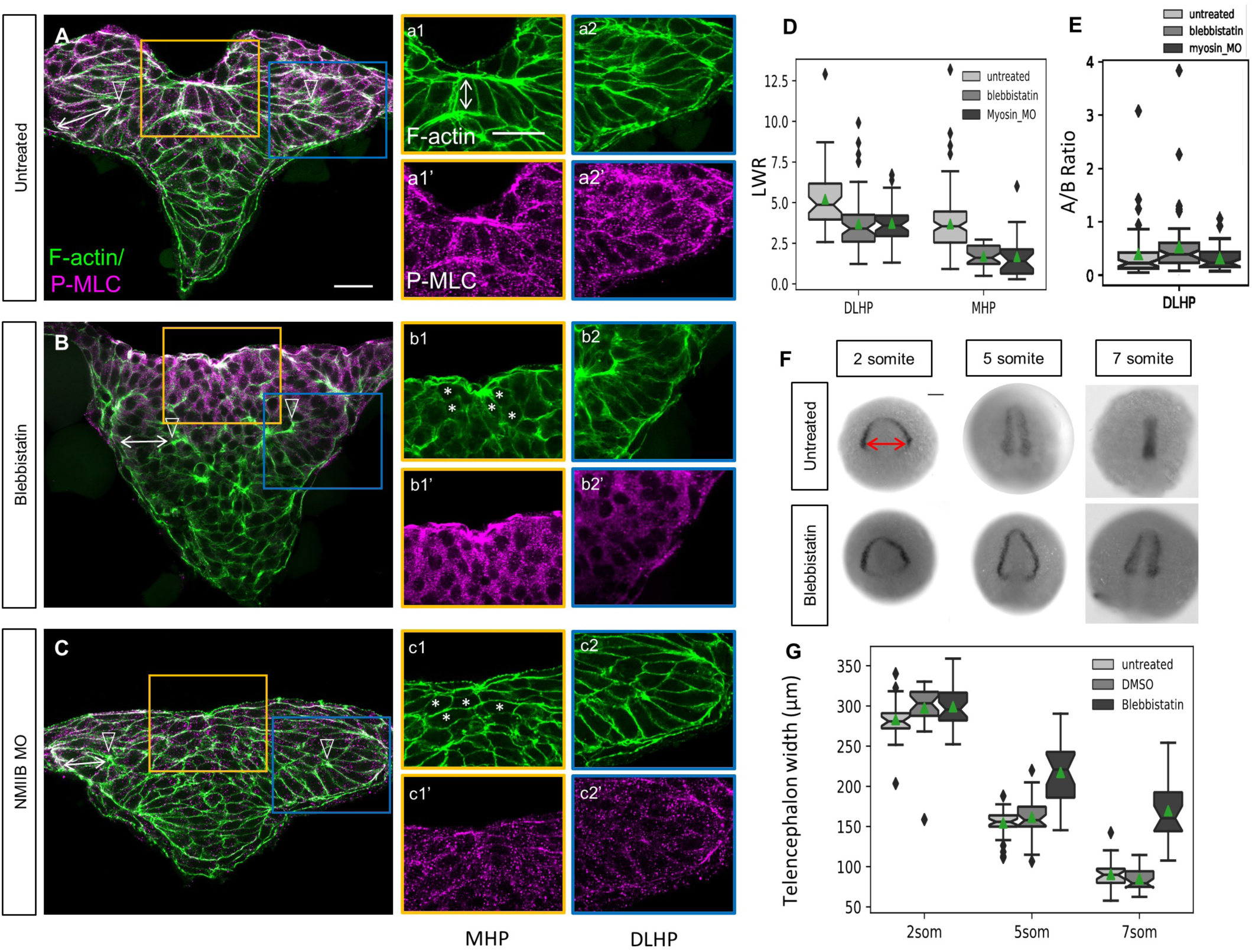
Role of non-muscle myosin II in apical constriction and neural fold convergence. Transverse sections through the ANP of 5 som untreated (A-a2’), blebbistatin-treated (B-b2’) and *NMIIB* MO-injected embryos (C-c2’). Embryos were labeled with phalloidin (F-actin, green) and anti-P-MLC (magenta). (D) Length-width ratio distributions for DLHP and MHP cells from untreated, blebbistatin-treated, and *NMIIB* MO-injected embryos. DLHP measurements: untreated: n = 50 cells (5 embryos), mean=5.22; blebbistatin-treated: n = 58 cells (4 embryos), mean=3.70; *NMIIB* MO-injected: n = 38 cells (3 embryos), mean=3.74. MHP measurements: untreated: n=115 cells (4 embryos), mean= 3.70 (same data as in Figure 5 c2); blebbistatin-treated: n= 35 cells (4 embryos), mean=1.70; *NMIIB* MO-injected: n= 24 cells (3 embryos), mean = 1.68. Two-sided Mann Whitney U test: DLHP: untreated vs blebbistatin: P=1.30e^-6; untreated vs *NMIIB* MO-injected: P = 2.14e^-5; MHP: untreated vs blebbistatin: P=3.55e^-12; untreated vs *NMIIB* MO-injected: P = 3.50e^-8. (E) Apical-to-basal length ratios of DLHP cells for untreated, blebbistatin-treated and *NMIIB* MO-injected embryos, same cells quantified as in D. Two-sided Mann Whitney U test: untreated vs blebbistatin-treated P= 0.00665; blebbistatin-treated vs *NMIIB* MO-injected: P=0.00827; untreated vs *NMIIB* MO-injected: P=0.930. (F) Dorsal views of 2, 5 and 7 som embryos untreated or blebbistatin-treated labeled via *in situ* hybridization using an *emx3* riboprobe. (G) Boxplots showing distribution of telencephalon widths (double red arrow in F) according to treatment group. Notches depict the 95% confidence interval around the median and green triangles depict distribution means. 2 som: Untreated: n=26, mean=283.224; DMSO-treated: n= 26, mean=297.727; blebbistatin-treated: n = 34, mean=299.412. 5 som: Untreated: n=33, mean=154.688; DMSO: n=30, mean=161.747; blebbistatin-treated: n=23, mean=217.472. Two-sided Mann Whitney U test: 5 som - untreated vs DMSO: P=0.332; untreated vs blebbistatin-treated: P=1.30e^-7; DMSO-treated vs blebbistatin-treated: P = 3.50e^-6. 7 som: untreated: n=28, mean=90.444; DMSO-treated: n=27, mean 84.855; blebbistatin-treated: n=24, mean= 169.779. Two-sided Mann Whitney U test: 7 som - untreated vs DMSO: P=0.170; untreated vs blebbistatin-treated: P=2.06e^-9; DMSO-treated vs blebbistatin-treated: P = 1.46e^-9. Annotations: double white arrows = cell length in deep layer; open arrowhead = DLHP; asterisks = rounded neuroectodermal cells; red double arrow = posterior-most telencephalon width. Scale bars: 25 μm in A and a1; 100 μm in B.

At a morphological level, disruption of the actomyosin network causes neural tube defects that trace back to impaired apical constriction and convergent extension in *Xenopus* embryos^15^. To test whether myosin is similarly required for forebrain neural tube closure in zebrafish, control (untreated and DMSO-treated) and blebbistatin-treated embryos were labeled at the 2, 5 and 7 som stage via *in situ* hybridization using the telencephalon marker *emx3* (Figure 10F) and the width of the posterior-most *emx3* domain was measured (Figure 10G). In contrast to control embryos, NF convergence was impaired in blebbistatin-treated embryos, beginning at the 5 som stage. It is possible that earlier convergent extension defects in the ANP contribute to this phenotype. However, failure of MHP cells to undergo apical constriction is likely to be a contributing cause given that disruption of several other proteins implicated in this process, including Shroom3^13^ and GEF-H1, a RhoA-specific GEF^14^ results in severe neural tube closure defects.

## DISCUSSION

We report here on mechanisms of forebrain morphogenesis in the zebrafish embryo and reveal that this region of the brain is formed by primary neurulation, involving the use of hingepoints and NFs.

The zebrafish MHP and paired DLHPs form in the superficial and deep layers of the eye field, respectively. The timing of MHP and DLHP formation overlap in zebrafish, in contrast to the biphasic nature of these events in mouse^3^. The zebrafish MHP is cup-shaped and more transient than its mammalian counterpart since these cells eventually intercalate between deep layer cells, contributing to the expansion of the eye vesicle^37^. Hingepoints are restricted to the ANP in zebrafish, however they are also present in more posterior regions in amniotes. Despite this difference, individual cells in the medial zone of the hindbrain neural plate were recently shown to internalize via a myosin-dependent mechanism^46^. Such variation from the organized cell clusters forming hingepoints in the zebrafish forebrain could be explained by precocious epithelialization of the ANP^37^. It thus appears that there is a transition from clustered internalization in the forebrain to individual cell internalization in the hindbrain region.

A key feature of hingepoint cells is their reduced apical surface, which is in part due to actomyosin contractility^13,41,47,48^. Apical constriction is thought to function as a purse string to generate the force required to bring the NFs together during cranial neurulation^49^. We provide evidence that the zebrafish MHP also utilizes an actomyosin-based contractile system. Assembly of this actomyosin network occurs via oscillatory constrictions with gradually decreasing amplitude, akin to the ratchet model initially proposed in invertebrates^50–52^ and later reported during neural tube closure in *Xenopus*^53^. We further show that disruption of myosin impairs NF convergence, possibly by contributing a “pulling force” on the NFs or by clearing the dorsal midline of eye field cells. These observations suggest that the actomyosin machinery is used across vertebrate to drive cranial neural tube closure and it will be interesting in the future to test whether upstream regulators of apical constriction such as Shroom3^13^ are also conserved.

In contrast to the MHP, the paired DLHPs do not require myosin to apically constrict, suggesting that DLHP formation is regulated by a distinct mechanism. Consistent with this observation, cell packing at a dorso-ventral boundary in the mouse neural tube causes buckling of the neuroectoderm at the DLHPs^54^. It is possible that such a mechanism also operates during zebrafish neurulation. Actomyosin may however generate cortical tension^15^, enabling both MHP and DLHP cells to maintain their elongated shape.

NFs in chick embryos form via epithelial ridging, kinking, delamination and apposition^16^, although the cellular basis of these morphogenetic events is not well understood. Elevation of the NFs in the mouse cranial neural plate is dependent on MHP constriction and expansion of the head mesenchyme^55^. NFs in the zebrafish are restricted to the ANP and head mesenchyme is unlikely to play a significant mechanical role in NF elevation, as the mesoderm layer underlying the ANP is thin. We instead identify basal constriction as a NF-intrinsic cell behavior that functions as a “reverse hingepoint”. Basal constriction may contribute to the early stages of NF elevation or ridging/kinking (implicated in the acquisition of the bilaminar topology of the NFs). Based on scanning EM images of chick embryos, it appears that this cell behavior may also be conserved in amniotes^16,17^.

The final step of primary neurulation involves the convergence and fusion of the NFs at the dorsal midline. NF convergence in zebrafish is likely to be mediated by cell intrinsic forces, including MHP and DLHP formation, although it is difficult to parse out their relative contribution, in addition to extrinsic forces derived from the non-neural ectoderm^17^. NF fusion is initiated at closure points in mammals and birds^56^. We observe two closure points in the zebrafish forebrain that form an eye-shaped opening that narrows from the corners in a bidirectional manner. Fusion of the NFs is mediated by the formation of cell protrusions that span the midline and establish the first points of contact with NF cells from the contralateral side. Akin to mice^21,22^, the cells that initiate contact between apposing NFs in zebrafish derive from the neuroectodermal portion of the NFs. However, rather than establishing contact andadhering via their lateral surfaces, the protrusive ends of zebrafish neuroectoderm cells appear to transiently interdigitate between their contralateral counterparts, forming a rod-like structure, the precursor of the telencephalon. These hyperextensions are later retracted as the telencephalon eventually epithelializes, forming a clearly defined midline. Once the neuroectoderm cells have met and fused, non-neural ectoderm cells complete their migration and fuse at the dorsal midline, forming a continuous epidermal layer.

In addition to the epidermis, frog and fish embryos are surrounded by an outer protective epithelial monolayer, called EVL in zebrafish. Given that the EVL is in direct contact with the NFs, it is possible that this epithelial layer contributes to forebrain morphogenesis by providing a stable substrate for NF convergence. Conversely, the EVL is transiently tugged inward as the MHP cells undergo apical constriction. It is therefore likely that the interactions between the neural ectoderm and the EVL are reciprocal.

Taken together, these findings reveal striking similarities and some unique features of forebrain neurulation in fish that have significant implications for our understanding of the evolution of neurulation and the relevance of zebrafish to understand human neural tube development.

## METHODS

### Zebrafish strains/husbandry

Studies were performed using wildtype (AB) strains or *Tg(emx3:YFP)*^*b1200* 39^ and embryos were raised at 28.5°C. All experiments were approved by the University of Maryland, Baltimore County’s Institutional Animal Care and Use Committee (IACUC) and were performed according to national regulatory standards.

### Nucleic acid and morpholino injections

Plasmids encoding membrane-targeted Green Fluorescent Protein (mGFP) (Richard Harland, University of California, Berkeley, CA, USA), membrane-targeted Red Fluorescent Protein (mRFP), membrane-targeted Kaede (mKaede) (a gift from Ajay Chtinis, National Institute of Health, Bethesda, MD) and *pard3:egfp*^57^ were linearized with NotI and transcribed using the SP6 mMESSAGE mMACHINE kit (Ambion, AM1340). For ubiquitous expression of *mGF*P, *mRFP, mkaede* or *par3d:eGFP*, 50 pg of RNA was injected into 1-cell stage embryos. For mosaic expression of mGFP, 50 pg of RNA was injected into 1 or 2 of the four central blastomeres at the 16-cell stage. These blastomeres have a high probability for neural fate^58^ and are easy to identify for reproducible injections.

MOs were designed and synthesized by GeneTools (Philomath, Oregon, USA) and injected into 1-cell stage embryos. *mhy10*(Non-muscle Myosin IIB, EXON2-intron2) was delivered at 3ng per injection.

*mhy10*: 5’-CTTCACAAATGTGGTCTTACCTTGA-3′ ^45^

Microinjections were performed using a PCI-100 microinjector (Harvard Apparatus, Holliston, MA, USA).

### Blebbistatin treatment

90% epiboly embryos were manually dechorionated in an agarose-covered petri dish with E3 medium. Once embryos reached the tailbud stage, they were placed into 50μM blebbistatin (B0560 Sigma-Aldrich) diluted with E3 and then incubated at 28.5°C. Stock solution of blebbistatin was prepared with DMSO as per manufacturer’s instructions. Accordingly, a control group of embryos were treated with 1% DMSO diluted with E3 alongside every blebbistatin trial. Once the desired developmental stage was reached (2-, 5-, or 7-som), embryos were immediately fixed with 4% paraformaldehyde (PFA). As blebbistatin is light sensitive, embryos were kept in the dark as much as possible until fixation.

### Fixed tissue preparations and immunolabeling

Embryos were fixed in 4% PFA for 16 hours at 4°C overnight. Immunolabeling was performed on whole mount embryos, which were then sectioned with a Vibratome (Vibratome 1500 sectioning system), with the exception of N-cadherin, which was done on sections. Primary antibody incubation was performed for 24-48 hours at 4°C and secondary antibody incubation for 2.5 hours at room temperature.

Antibodies used: Rabbit anti-GFP at 1:1000 (Invitrogen, A11122), Rabbit anti-Sox3c at 1:2000 (Gift from Michael Klymkowsky), Rabbit anti-P-myosin light chain at 1:50 (Cell Signaling Technology, #3671S), Mouse anti-p63 at 1:200 (Santa Cruz BioTechnology, SC-8431 no longer in production), Rabbit anti-N-cadherin at 1:50 (Abcam, ab211116) and Mouse anti-ZO-1 antibody at 1:100 (Invitrogen, 33-9100). Alexa Fluorophore secondary antibodies were all used at a 1:1000 concentration: Goat anti-Rabbit −488, −568, −594 and Goat anti-Mouse −488, −594.Alexa Fluor 488-conjugated or 594-conjugated Phalloidin (Invitrogen, A12379 and A12381) at 1:250 and DAPI (Invitrogen, D1306) were used according to manufacturer’s instructions. Sections were mounted on glass slides using ProLong Diamond Antifade Mountant (Invitrogen, P36961). *Tg(emx3:YFP)*^*b1200*^ embryos were immunolabeled with anti-GFP to amplify the signal. mGFP and *pard3:eGFP*-injected embryos were not immunolabeled with anti-GFP.

### Whole-mount *in situ* hybridization and imaging

*In situ* hybridization was performed as described^59^. *emx3* riboprobe template was generated by PCR amplification using cDNA from 24hpf embryos.

T7 promoter: **TAATACGACTCACTATAGGG**

*emx3* antisense:

FWD: TCCATCCATCCTTCCCCCTT

RVS: **TAATACGACTCACTATAGGG**GTGCTGACTGCCTTTCCTCT

DIG-labeled riboprobes were generated using 2ul of PCR template with the Roche DIG RNA Labeling Kit (T7) (Sigma aldrich, SKU 11277073910)

Whole-mount imaging was carried out using a Zeiss Axioscope2 microscope. Embryos were imaged in a 2.5% glycerol solution.

### Confocal microscopy

#### Dorsal view time-lapse microscopy

performed as previously described^60^. Embryos were imaged using a Leica confocal microscope (Leica SP5 TCS 4D) at 15 sec/frame capturing <.5μm of tissue. All fluorescently labeled sections were imaged using a Leica confocal microscope (Leica SP5 TCS 4D).

#### Transverse view time-lapse microscopy

performed as described previously^60^, with the exception that the embryos were oriented anterior side against the glass. Embryos were imaged using a Zeiss confocal microscope (Zeiss 900 LSM with Airyscan 2), at 20X.

### Data Quantification

Medial-most-p63-positive-domain migration, Figure 1N: For each hemisphere of a tissue section, the distance between the medial-most p63 positive nucleus and the midline was manually scored. For each embryo, measurements were taken from tissue sections ranging along the anterior-posterior axis of the forebrain.

Deep layer cell morphology measurements, Figure 3: Cells were manually scored from non-projected z-stack images, where cellular outlines were visually determined from the mGFP signal. Cells were labeled as neuroectoderm NF cells based on the fan-like pattern of their basal projections in contact with the non-neural ectoderm. All other cells scored were within the morphologically distinct eye vesicle and were labeled as eye field cells. Neural ectoderm NF cells and eye field cells are not morphologically distinguishable at the 2 somitic stage and thus were not given cell type identities at that developmental stage.

Medial hinge cell morphology measurements, Figure 5C: Cells were manually scored from non-projected z-stack images of the overlay between the labeled actin cytoskeleton (phalloidin) and nucleus (DAPI, not shown in Figure 5) channels. Cellular outlines were visually determined from the phalloidin signal. For the nuclear position measurement (Figure 5c3), the distance from the dorsal edge of the nucleus to the basal membrane of the cell was scored and divided by the total cell length.

Cell ratcheting, Figure 6 a1,a2,b1,b2: Live movie z-stacks for the first 35 minutes (∼2-4 somites, n=2) of each movie were max projected and cropped so the midline of the tissue horizontally bisected the image frame. The mGFP signal was then inverted and thresholded to produce a binary image. Measurements for individual cell surface areas were captured using the magic wand tool in ImageJ for every frame (15 seconds) until the cell left the field of view. Plotted on the x-axis is the order of frame measurements (frame 0, frame 1, frame 2,…) divided by the total number of frames for which that cell was scored to standardize between 0 and 1. Plotted on the y-axis, the initial value of each cell’s surface area was subtracted from all measurements for that cell to initialize surface area’s to zero. The data was binned (bins=50) to obtain median measurements and confidence intervals. Cells that underwent mitosis anytime during the movie were excluded. Cells were labeled as MHP if their surface area centroid from the first frame (t=0) was +-20 μm from the midline with MHP-adjacent cells being labeled as such if their centroid was > +-20 μm from the midline.

#### Dorsal view movies of apical surface area oscillations

An oscillation was defined as a cycle of constriction and then expansion or vice versa. For a given cell, constriction or expansion was determined by the difference between surface areas for two sequential time frames, where frame_1_surface_area - frame_2_surface_area > 0 was scored as constricting and < 0 was scored as expanding.

#### Transverse view movies of cell shape changes

Movies were manually scored at each time frame to measure the following parameters (n=2, data shown for Supplemental movie 1 only). Neural groove formation was calculated as the angle formed by the dorsal-most medial tissue (Figure 8G); optic vesicle angle was measured as the angle formed by the outline of the optic vesicle as it forms (Figure 8H); NF basal angle was calculated as the angle formed by the basal surface of NF cells (Figure 8I); NF elevation was measured as the distance between the basal side of NF cells and the apical side of MHP cells (Figure 8J). As the NFs elevate, that distance decreases and eventually becomes negative as the NFs elevate above the MHP. Finally, the distance between the NFs was measured as the distance between the basal side of NF cells, as they migrate towards the midline (Figure 8K).

*emx3 in situ* hybridization measurements, Figure 10C: The distance between the lateral edges of the posterior-most extent of the emx3 domain was manually scored for each embryo.

### Statistical Analysis

The Mann-Whitney U test was used for all significance testing. The python function scipy.stats.mannwhitneyu(alternative=‘two-sided’) was used to calculate the test statistic and P-value for each significance test. Graphs were generated using the python Seaborn package with the following functions: seaborn.boxplot(), seaborn.lmplot(), seaborn.violinplot(), seaborn.lineplot().

### Data availability

The authors declare that all data supporting the findings of this study are available within the article and its supplementary information files or from the corresponding author upon reasonable request.

## ACKNOWLEDGEMENTS

Funds from Howard Hughes Medical Institute through the UMBC Precollege and Undergraduate Science Education Program supported J. Werner and D. Brooks. Funds from NIH/NIGMS grants # T32-GM055036 and # R25-GM066706 and NSF LSAMP BD grant # 1500511 to UMBC supported M. Negesse. Funds from NSF LSAMP grant # 1619676 to UMBC supported J. Johnson. Funds from NIH/NIGMS MARCU*STAR T34 grant # HHS 00026 to UMBC supported D. Brooks and A. Caldwell. We thank the following people for their contributions: Tagide deCarvalho for her help with confocal imaging and image processing; Corinne Houart for the *Tg(emx3:YFP)*^*b1200*^ transgenic line; Jennifer Gutzman for the gift of *myh10* morpholino and Mark Van Doren for his comments on the manuscript.

## AUTHOR CONTRIBUTIONS

J.W. and M.N. contributed equally to this study.

J.W. designed and performed experiments and carried out data analysis. M.N. carried out experiments and data analysis, annotated movie files and generated illustrations. D.B. and A.C. contributed to the experiments and analysis of cell polarity and myosin function. J.J. contributed to the analysis of myosin function. R.B. oversaw experimental design and analysis and wrote the manuscript.

## COMPETING INTERESTS

The authors declare no competing interests

## SUPPLEMENTARY MOVIE LEGENDS

**Supplemental Movie 1. MHP cells sink inwards as neural folds elevate.** Time lapse imaging of embryo ubiquitously expressing m Kaede (green) that was photo-converted in medial superficial cells that form the MHP (magenta). Annotations: Cyan arrowhead: basal surface of neural fold cells; white arrows: sinking of MHP cells.

**Supplemental Movie 2. 3-Dimensional rotation revealing organization of neural fold cells in a 5 som embryo.** Transverse section, at the level of the forebrain of a 5 somite embryo mosaically-expressing mGFP.

**Supplemental Movie 3. 3-Dimensional rotation revealing organization of neural fold cells in a 7 som embryo.** Transverse section, at the level of the forebrain of a 7 somite embryo mosaically-expressing mGFP.

**Supplemental Movie 4. Neural fold cells constrict basally as neural folds elevate.** Time lapse imaging of Tg[*emx3:YFP*] transgenic embryo ubiquitously expressing mRFP. YFP-positive neural fold cells (green) constrict basally as neural folds elevate. Annotations: Cyan arrow: basal surface of neural fold cells; white lines: telencephalon cells expressing YFP.

**Supplemental Movie 5. MHP cells undergo oscillatory constriction with decreasing amplitude.** Time lapse imaging of embryo ubiquitously expressing mGFP. Clusters of medial (MHP) constrict apically in an oscillatory manner, in contrast to their lateral neighbors that do not. Annotations: cyan dots = MHP cells; yellow asterisks = MHP-adjacent cells that do not undergo apical constriction, yellow dashed line: midline, red asterisks = EVL cells.

**Supplemental Movie 6. Neural tube closure is initiated at two closure points in the forebrain.** Time lapse imaging of embryo mosaically-expressing mGFP. The Green and brightfield channels are overlaid to reveal the shape of the neural folds and neural groove. Annotations: red asterisk: apex of the arch shaped neural folds; white dotted line: contour of the eye-shaped opening whose corners are defined by closure points 1 and 2, double headed arrow: width of the neural plate, which decreases over time, white arrows: closure points 1 and 2, respectively.

**Supplemental Movie 7. Neural fold cells extend filopodial protrusions across the midline.** Time lapse imaging of embryo ubiquitously expressing mGFP. Cells (cyan and magenta) originating from contralateral sides of the ANP extend medially oriented filopodia and transiently interdigitate across the midline (yellow dashed line).

## SUPPLEMENTAL FIGURE

**Supplemental Figure S1.**
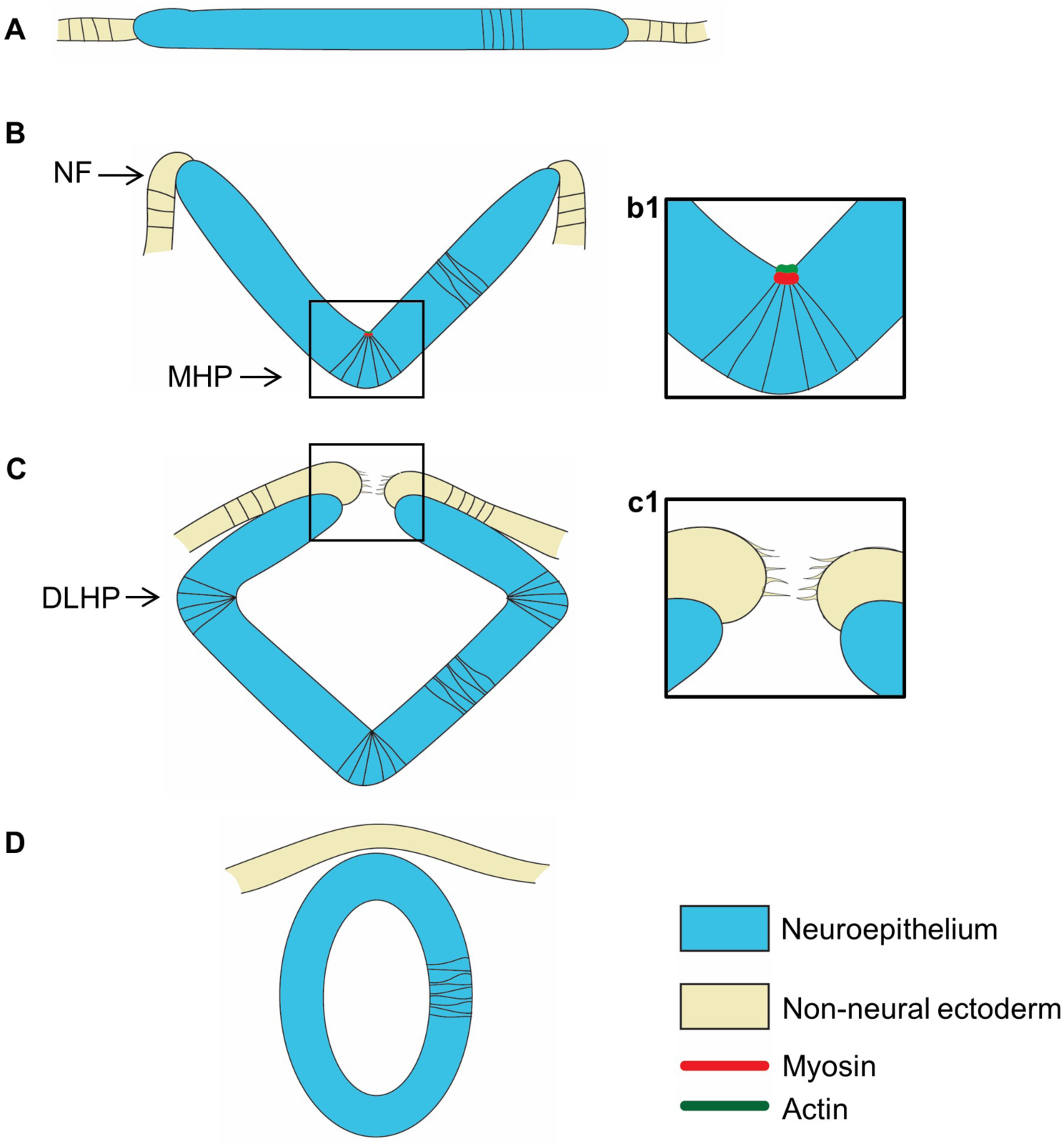
Neurulation in amniotes. Cross sectional illustration of stages of neurulation in amniotes. (A) The neural plate and adjacent non neural ectoderm. (B) Medial hingepoint formation shapes the neural groove and elevates the neural folds. (b1) Illustration of medial hingepoint cells that are apically constricted and enriched for actomyosin at their apex. (C) Dorso-lateral hingepoint formation brings the neural folds in close apposition. (c1) Filopodial extensions establish contact between neural fold cells across the midline. In the mouse forebrain the first contact is established between neuroectodermal cells. (D) The neural folds fuse medially, separating the epidermis from the neural tube. Abbreviations: DLHP = dorso-lateral hingepoint; MHP = medial hingepoint; NF = neural fold.

## Notes

### Competing Interest Statement

The authors have declared no competing interest.

