## Supplementary figures and images for "Hingepoints and neural folds reveal conserved features of primary neurulation in the zebrafish forebrain"

### Supplemental Figure 1

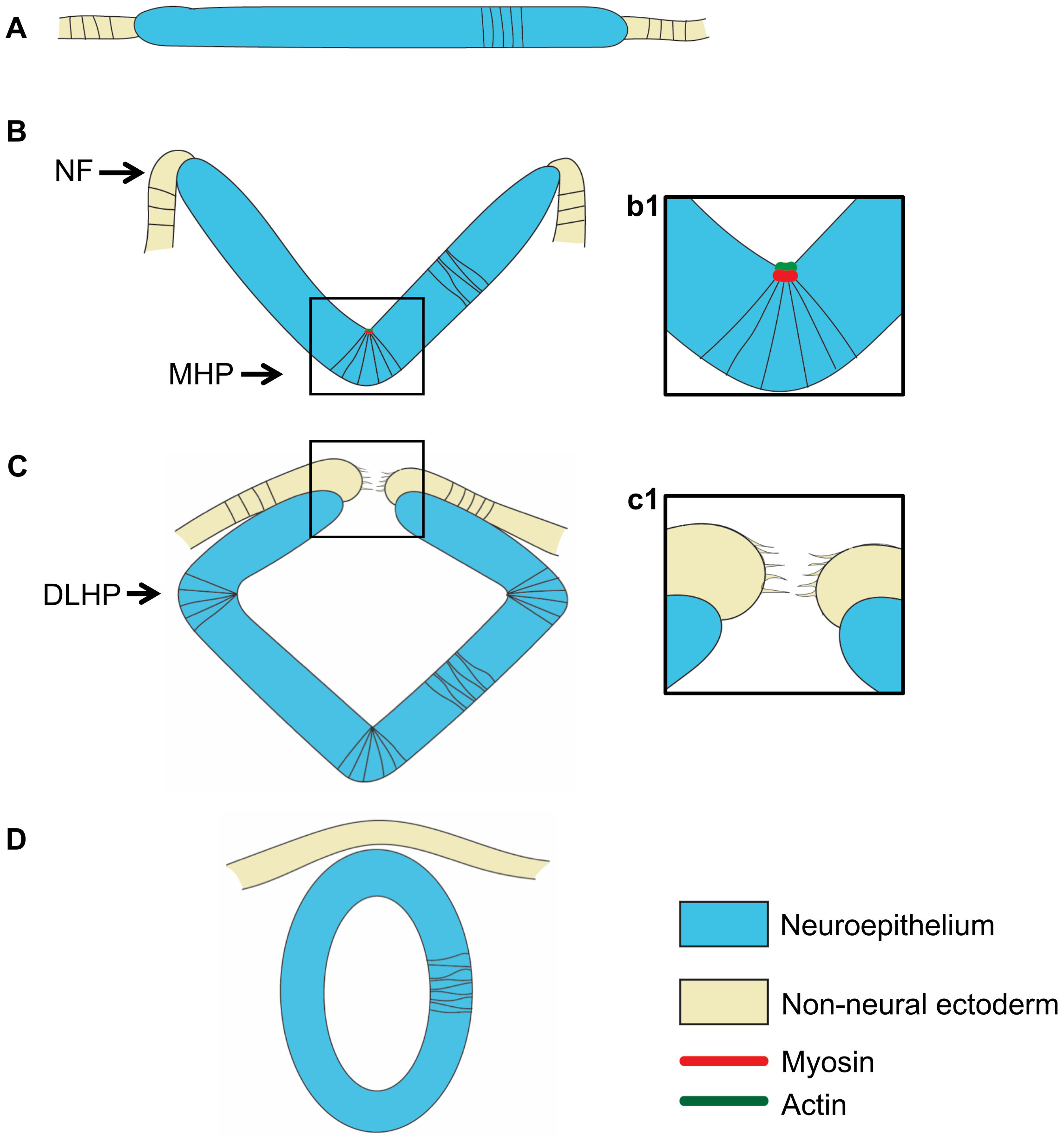
